# Myocardial STIM1 deficiency triggers mitochondrial fission to impair electrophysiological function and exacerbate post-MI remodeling

**DOI:** 10.1101/2024.12.05.625436

**Authors:** Marine Cacheux, Juan Velasco, Xiaohong Wu, Jonathan M. Granger, Benjamin Strauss, Fadi G. Akar

## Abstract

**Background:** Stromal interaction molecule 1 (STIM1) is a key regulator of intracellular calcium (Ca²⁺) homeostasis and mitochondrial function, two tightly coupled processes that critically determine electro-mechanical function and post-myocardial infarction (MI) remodeling.

**Hypothesis:** We hypothesized that cardiomyocyte STIM1 deficiency induced by AAV9-mediated knockdown triggers mitochondrial and metabolic remodeling that alters the electrophysiological (EP) substrate and exacerbates post-MI heart failure.

**Methods:** Eight-week-old mice received cardiotropic AAV9-mediated delivery of short hairpin RNA targeting STIM1 (AAV9-shSTIM1) or a control construct (AAV9-shLuc/PBS). Four weeks later, *in vivo* echocardiography followed by *ex vivo* optical action potential (AP) mapping were performed to determine the impact of myocardial STIM1 downregulation on mechanical and EP properties. Electron microscopy and western blotting were performed to assess mitochondrial network architecture, dynamics and related metabolic signaling. A separate cohort of mice underwent a 30-minute coronary occlusion followed by reperfusion to induce MI. One week post-MI, left ventricular (LV) function, structural remodeling, and EP properties were assessed in mice with and without myocardial STIM1 knockdown.

**Results:** AAV9-shSTIM1 delivery markedly reduced cardiomyocyte STIM1 expression. STIM1 downregulation induced extensive mitochondrial fragmentation, increased DRP1 phosphorylation at S616 (+55%, *p*=0.0057), reduced OPA1 expression, and altered AMPK-dependent signaling (reduced phospho-ACC and phospho-Raptor). These mitochondrial and metabolic abnormalities were accompanied by EP instability. In response to *in vivo* I/R injury, shSTIM1 mice exhibited significantly greater LV dysfunction (−44.3% vs. −12.2% fractional shortening, *p*<0.0001) and more pronounced structural and EP remodeling with formation of spatially discordant AP alternans 1-week following MI indicative of accelerated heart failure progression.

**Conclusions:** AAV9-mediated STIM1 knockdown exacerbates post-MI cardiac dysfunction likely through impaired mitochondrial dynamics and disrupted metabolic signaling. Beyond worsening post-MI function, STIM1 deficiency alone induces profound EP remodeling, underscoring the pivotal role of cardiomyocyte STIM1 in coordinating mitochondrial homeostasis, electrical stability, and cardiac recovery after injury.

## INTRODUCTION

STIM1 is a single-pass transmembrane protein that functions as a luminal calcium (Ca²⁺) sensor within the endoplasmic reticulum (ER).^1, 2^ Upon ER Ca²⁺ depletion, STIM1 undergoes conformational activation and engages its primary plasma membrane partner, Orai1, to initiate the canonical process of store-operated Ca²⁺ entry (SOCE), facilitating Ca²⁺ influx from the extracellular space. Although initially characterized in non-excitable cells, STIM1-dependent SOCE has since been identified in a host of excitable cell types, including skeletal^3^ and smooth muscle cells,^4^ neurons,^5^ and cardiomyocytes.^6, 7^

In the heart, STIM1 expression is dynamically regulated by developmental stage and disease state. It is abundant in neonatal cardiomyocytes, where SOCE drives Ca²⁺-dependent signaling required for growth and maturation.^6^ In contrast, adult cardiomyocytes exhibit markedly reduced STIM1 expression and rely instead on the classical excitation–contraction coupling machinery (L-type Ca²⁺ channels, SERCA2a, phospholamban, ryanodine receptor type 2, Na⁺– Ca²⁺ exchanger, etc), to support rapid, beat-to-beat Ca²⁺ cycling. Pathological stimuli such as pressure overload re-induce STIM1 expression, augmenting SOCE and promoting cardiac hypertrophy *in vitro* and *in vivo*^8, 9^. Cardiac-specific STIM1 overexpression in transgenic mice recapitulates this phenotype, leading to hypertrophy and heart failure, effects prevented by pharmacological SOCE inhibition.^7, 10, 11^ Notably, SOCE acts as a slow, adaptive mechanism that replenishes sarcoplasmic reticulum (SR) Ca²⁺ stores rather than modulating beat-to-beat Ca²⁺ transients.

Beyond its role in mediating SOCE, Zhao et al.^12^ demonstrated that STIM1 overexpression in rat ventricular myocytes increases SR Ca²⁺ content through direct interaction with phospholamban highlighting its capacity to influence Ca²⁺ handling and contractile function on a beat-by-beat basis. Consistent with this, our prior work using a tamoxifen inducible, cardiomyocyte-specific STIM1 knockout model (αMHC-MerCreMer system) revealed that acute STIM1 depletion (>80% reduction) provokes electrophysiological (EP) instability characterized by Ca²⁺-mediated afterdepolarizations, spatially discordant action potential (AP) alternans, and sustained ventricular fibrillation.^13^ However, these EP abnormalities emerged in the context of rapid mortality and severe left ventricular (LV) dysfunction, precluding definitive mechanistic attribution to primary STIM1 loss *per se*. The underlying mechanisms linking STIM1 depletion to these EP defects therefore remain incompletely understood.

Emerging evidence extends the functional repertoire of STIM1 beyond Ca²⁺ signaling to encompass mitochondrial regulation and cellular metabolism.^14–17^ STIM1 physically interacts with mitofusin-2 (MFN2), a key mitochondrial dynamics protein, maintaining mitochondrial morphology and SR–mitochondrial tethering.^18, 19^ Genetic ablation of STIM1 alters substrate utilization, underscoring its essential role in metabolic homeostasis. Furthermore, STIM1 is phosphorylated by the cellular energy sensor AMPK at specific serine residues that modulate SOCE in response to stress and exercise,^20, 21^ establishing a bidirectional coupling between metabolic signaling and Ca²⁺ homeostasis within cardiomyocytes.

Given STIM1’s dual role in Ca²⁺ and metabolic regulation, we hypothesized that cardiomyocyte STIM1 downregulation alters electromechanical function by disrupting mitochondrial and metabolic processes. We further postulated that these changes would sensitize the myocardium to acute myocardial infarction (MI) following *in vivo* ischemia/reperfusion (I/R) injury, a condition characterized by Ca²⁺ overload and mitochondrial dysfunction. Whereas smooth muscle–specific STIM1 deletion confers protection against MI^22^, we anticipated that loss of STIM1 in cardiomyocytes would have the opposite effect, impairing Ca²⁺ and metabolic signaling and exacerbating post-MI remodeling.

To test this dual hypothesis, we employed a cardiotropic adeno-associated virus serotype 9 (AAV9) vector encoding a short hairpin RNA targeting STIM1 (AAV9-shSTIM1), a strategy validated for efficient myocardial STIM1 knockdown (**Data Supplement 1**).^11^ Using this approach, we first assessed cardiac mechanical and EP function four weeks following gene transfer and examined associated mitochondrial and metabolic alterations. In a separate cohort, we assessed post-MI function and remodeling in mice with and without AAV9-mediated STIM1 knockdown.

Our findings demonstrate that myocardial STIM1 downregulation was associated with moderate LV dysfunction 4 weeks following gene transfer. This was accompanied by abnormal EP behavior, including conduction slowing, susceptibility to pacing-induced AP alternans and irregular rhythm. Mechanistically, STIM1 loss induced excessive mitochondrial fission, yielding a fragmented network of smaller organelles and suppressed AMPK-dependent downstream metabolic signaling. Following MI, hearts with STIM1 downregulation exhibited accelerated progression to end-stage heart failure with severe LV dysfunction and a more advanced EP remodeling phenotype characterized by the emergence of spatially-discordant alternans.

## METHODS

Primary data supporting the findings of this study are available from the corresponding author upon reasonable request. All experimental procedures complied with the *National Institutes of Health* Guide for the Care and Use of Laboratory Animals. Protocols were approved by the Institutional Animal Care and Use Committee.

### AAV9-mediated in-vivo gene transfer

Recombinant AAV9.sh*Stim1* and AAV9.sh*Luc* (against luciferase) were produced as previously described.^11^ Eight-week old C57bl/6J male mice received 1×10^11^ viral genomes (vg) of AAV9.sh*Stim1* (**Data Supplement 1**) or AAV9.sh*Luc* or PBS by tail vein injection.

### In-vivo model of I/R injury to induce MI

Four weeks following gene transfer or PBS injection, mice underwent left coronary artery (LCA) ligation for 30 minutes followed by reperfusion to induce MI (**Figure 2A**). For this procedure, anesthesia was induced via a single intraperitoneal (IP) injection of a mixture containing ketamine (1 mL × 100 mg/mL), xylazine (0.1 mL x 20 mg/mL), acepromazine (0.1 mL × 10 mg/mL), in 0.9% normal saline (1 mL final). Once anesthetized, the left side of the chest was shaved. The mouse was intubated with a 20G catheter and ventilated. Body temperature was maintained using a heating pad. The animal’s right hind paw was secured to the surgical table, with the left hind paw taped over the right, positioning the left side of the chest upwards. The skin was disinfected with 10% povidone-iodine. An incision was made at the fourth intercostal space to access the chest, followed by opening the pericardium. The LCA was temporarily ligated at the mid-level using a 7.0 silk suture. Successful ligation was confirmed by observing tissue pallor. After 30 minutes, the ligature was released, and reperfusion was verified by the return of the apex to its normal color. The chest was closed using 6.0 silk suture. Buprenorphine (0.05 mg/kg) was administered via IP injection for analgesia. Ventilation was discontinued once spontaneous breathing resumed, and the mouse was placed on a heating pad until fully conscious. Postoperative monitoring continued daily for one week, with additional buprenorphine injections (0.05 mg/kg) provided every 12 hours for 48 hours to manage pain.

### Echocardiography

Non-invasive echocardiography was performed under sedation by IP injection of up to 75mg/kg of ketamine to allow adequate restraint of the animal while maintaining a heart rate of ∼600 beats per minute. The animal chest was shaved. Left ventricular (LV) walls thickness and internal diameter were measured by M-mode using a Vivid 7 echocardiography apparatus with a 13-MHz linear array probe (General Electric, New York, NY). Short-axis parasternal views of the LV at the midpapillary level were used to guide the M-mode measurements. At least three different measurements of the intraventricular septum (IVS) and LV posterior wall (LVPW) dimensions as well as the LV internal diameter (LVID) were averaged per animal in systole (s) and diastole (d). LV performance was assessed by fraction shortening (FS) measurement as follows: FS = (LVIDd – LVIDs) / LVIDd. Videos of 2D transthoracic parasternal long axis view echocardiography were also recorded (**Data Supplement 2**). Echocardiography measurements were performed at 3 timepoints: baseline prior to gene transfer (8 weeks of age), 4 weeks post gene transfer (12 weeks of age) and 1 week later which also corresponds to 1 week post MI surgery (13 weeks of age).

### Optical AP mapping of ex-vivo perfused hearts

Hearts were excised rapidly and perfused on a Langendorff apparatus with Tyrode’s solution containing (in mM): NaCl 129, NaHCO_3_ 24, KCL 4, MgCl_2_ 1.0, Glucose 11, CaCl_2_ 1.8, KH_2_PO_4_ 1.2, pH 7.4). Perfusion pressure was maintained at ∼70 mmHg by regulating coronary perfusion flow throughout the experiment. Preparations were positioned in a tissue bath gently pressed against a glass imaging window as previously described.^23–25^ Movement was minimized by perfusion with blebbistatin. To avoid surface cooling, preparations were immersed in the coronary effluent maintained at a physiological temperature by a heat exchanger assembly. Hearts were stimulated with a pacing electrode placed on the anterior epicardial surface of the LV, mid-way between apex and base. Cardiac rhythm was monitored continuously with silver electrodes connected to an electrocardiogram amplifier. Optical APs were measured with an 80×80-pixel CCD camera as previously described.^23–25^ High-resolution optical APs were measured simultaneously from 6,400 sites at 1ms temporal resolution during each recording. To improve signal quality, spatial binning was performed, which yielded an array of 400 (20×20) high-fidelity optical APs that were amenable to accurate automated analyses. The experimental protocol consisted of pacing hearts at progressively shorter pacing cycle lengths (PCL) starting at 140ms and in decrements of 10ms up to PCL 50ms. At each PCL, the stimulus train had a duration of at least 1min. Steady-state APD and CV values were determined as previously described. ^23^ ^24, 25^ Repolarization times at 75% relative to the AP amplitude were quantified and APD was defined as the temporal difference between the repolarization and activation times at each site. Local activation time at each site was defined as the maximum first derivative during the upstroke of the AP. Velocity vectors (magnitude and direction) were derived from the activation times of each pixel relative to those of its neighbors. CV was measured by averaging the magnitude of the velocity vectors.

### Western blots

15 or 30ug of protein lysates were loaded on 4-15% tris-glycine gels (Biorad). Primary antibodies used were STIM1 (Sigma-Aldrich 6197), GAPDH (Cell Signaling 2118), DRP1 (Cell Signaling 8570), pDRP1-S616 (Cell Signaling 3455), pDRP1-S637 (Cell Signaling 4867), MFN1 (Proteintech 13798-1-AP), MFN2 (D2010) (Cell Signaling 9482), OPA1 (D6UN6) (Cell Signaling 80471), tAMPK (Cell Signaling 2532), tACC (Cell Signaling 3661), pACC (Cell Signaling 3661), tRaptor (Cell Signaling 2280), pRaptor-S792 (Cell Signaling 2083), Cx43 (Cell Signaling 3512) and Nav1.5 (Alomone #ASC-005) were used. Secondary antibodies were species-specific horseradish peroxidase-conjugated antibodies.

### Electron microscopy

Murine hearts were rapidly excised and cannulated through the aorta. Hearts were perfused for 3 min with PBS at 37°C to wash out the blood followed by a fixative solution of Glutaraldehyde 3% in 0.1M Cacodylate buffer pH7.4 (Electron Microscopy Sciences-cat#16538-15) for less than 5 min. The LV was then dissected in samples of 1mm x 2mm in dimension. LV samples were placed into the fixative buffer for 2h at room temperature and kept overnight at 4°C. Samples were processed by the EM Core facility. EM Images were quantified using *Image J*.

### Statistical Analyses

Unpaired Student *t* test was used to compare differences between 2 groups. For multiple comparisons, two-way ANOVA followed by Sidak’s multiple comparisons test was used. P<0.05 was considered significant.

## RESULTS

We employed a cardiotropic AAV9-mediated gene transfer strategy to achieve myocardial knockdown of STIM1 levels using a short hairpin RNA construct (AAV9-shSTIM1) (**Figure 1A**). Four weeks after gene delivery, cardiomyocyte STIM1 expression was markedly reduced in shSTIM1 hearts (**Figure 1B**). Importantly, this reduction alone was sufficient to impair mechanical function, as evidenced by a moderate decrease in fractional shortening on echocardiography (42.3 ± 1.7% in shSTIM1 vs. 59.9 ± 1.2% in Ctrl, *p* < 0.001) (**Figure 1C&D, Data Supplement 2)**. This functional decline was accompanied by mild LV dilation, reflected by an increase in the LV internal diameter (LVIDd: 3.32 ± 0.1 mm in shSTIM1 vs. 2.85 ± 0.05 mm in Ctrl, *p* = 0.0002), without significant changes in interventricular septal thickness (IVSd), posterior wall thickness (LVPWd), or heart rate (**Figure 1E**).

**FIGURE 1:**
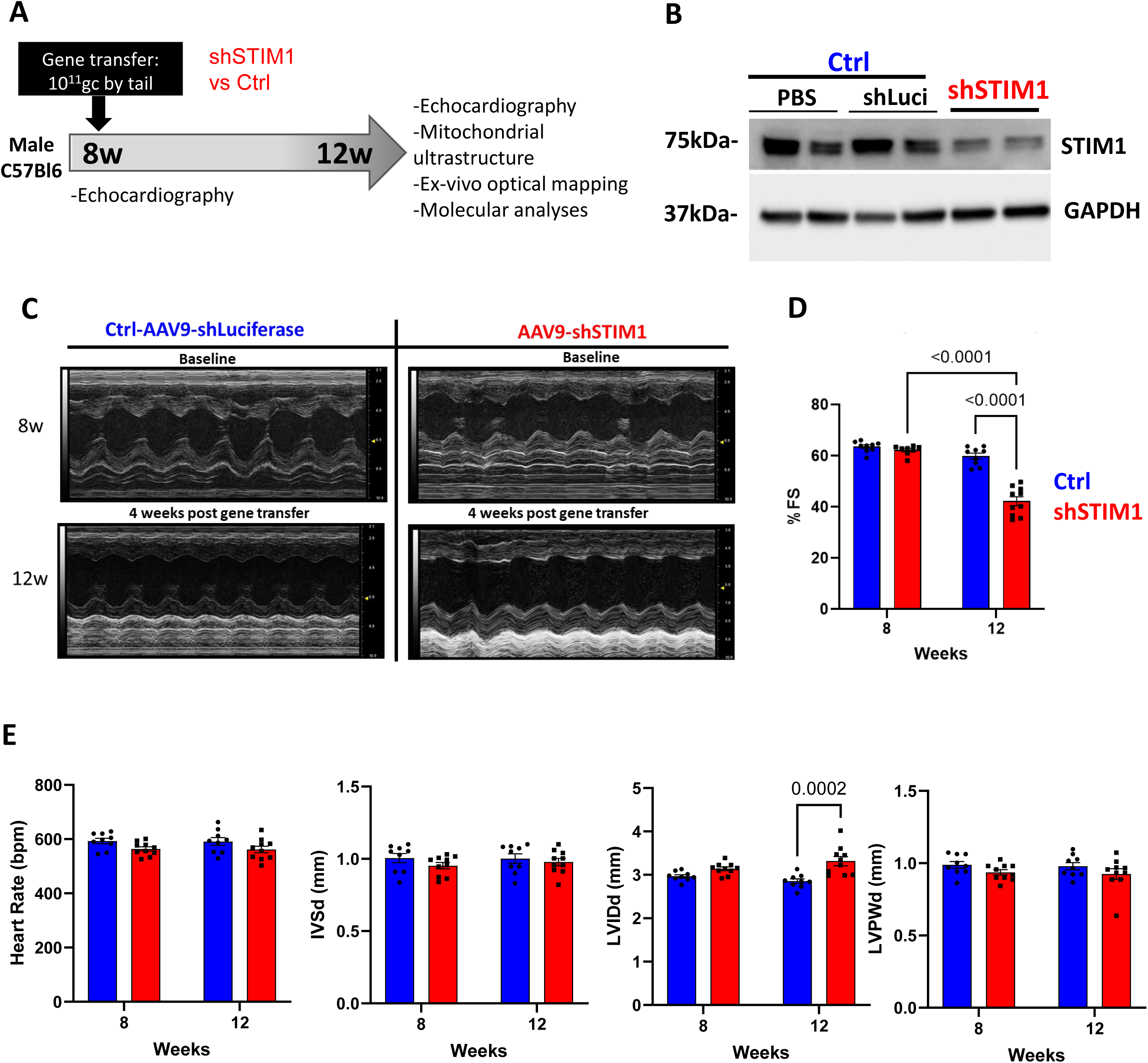
AAV9-mediated STIM1 silencing induced mechanical and structural LV remodeling. **A.** Schematic illustrating the experimental timeline to assess the impact of cardiomyocyte STIM1 downregulation per se in altering mitochondrial ultrastructure, EP properties and LV function. After baseline echocardiographic measurements, 8 weeks old C57Bl6/J male mice underwent AAV9-mediated shSTIM1 gene transfer via a tail vein injection with 1×10^11^ vg. Control (Ctrl) mice received either an AAV9 carrying a short hairpin against luciferase (shLuci) or PBS injection. Four weeks post gene transfer (at 12 weeks of age), mice underwent echocardiographic measurements before experimental procedures. **B.** Representative western blot documenting efficiency of STIM1 downregulation in lysates from isolated cardiomyocytes of shSTIM1 versus control (PBS and shLuci injected hearts). Cardiomyocyte STIM1 expression is reduced relative to GAPDH in shSTIM1 hearts 4 weeks post gene transfer. **C.** Representative echocardiographic M-mode images at baseline (8w old) and 4 weeks post gene transfer (12w old) in Ctrl and shSTIM1 mice. Echocardiographic parameters measured were: (**D**) fractional shortening (%FS), (**E**) heart rate, interventricular septum thickness (IVSd), LV internal diameter (LVIDd) and LV posterior wall thickness (LVPWd) during diastole. N=10 AAV9.shSTIM1 mice and N=9 control mice were used for this analysis. Statistics: two-ways-ANOVA between Ctrl and shSTIM1, with SD error bars.

We next examined EP properties in AAV9-shSTIM1 and Ctrl mice 4 weeks following gene or sham delivery, respectively (**Figure 2)**. While mean APD₇₅ values did not differ between groups (**Figure 2A**), shSTIM1 hearts exhibited significant conduction velocity (CV) slowing (40.7 ± 3.0 cm/s vs. 51.2 ± 3.1 cm/s in Ctrl, *p* = 0.04), as illustrated by the representative isochrone maps (**Figure 2B–C**). This conduction defect was not attributable to sodium channel (Nav1.5) expression, which was paradoxically upregulated by 36% in shSTIM1 hearts (135.6 ± 11.7% vs. 100 ± 9.3% in Ctrl). Instead, it coincided with a 22% reduction in connexin 43 (Cx43) expression (77.7 ± 3.5% vs. 100 ± 8.4% in Ctrl) (**Figure 2D-E**).

**FIGURE 2:**
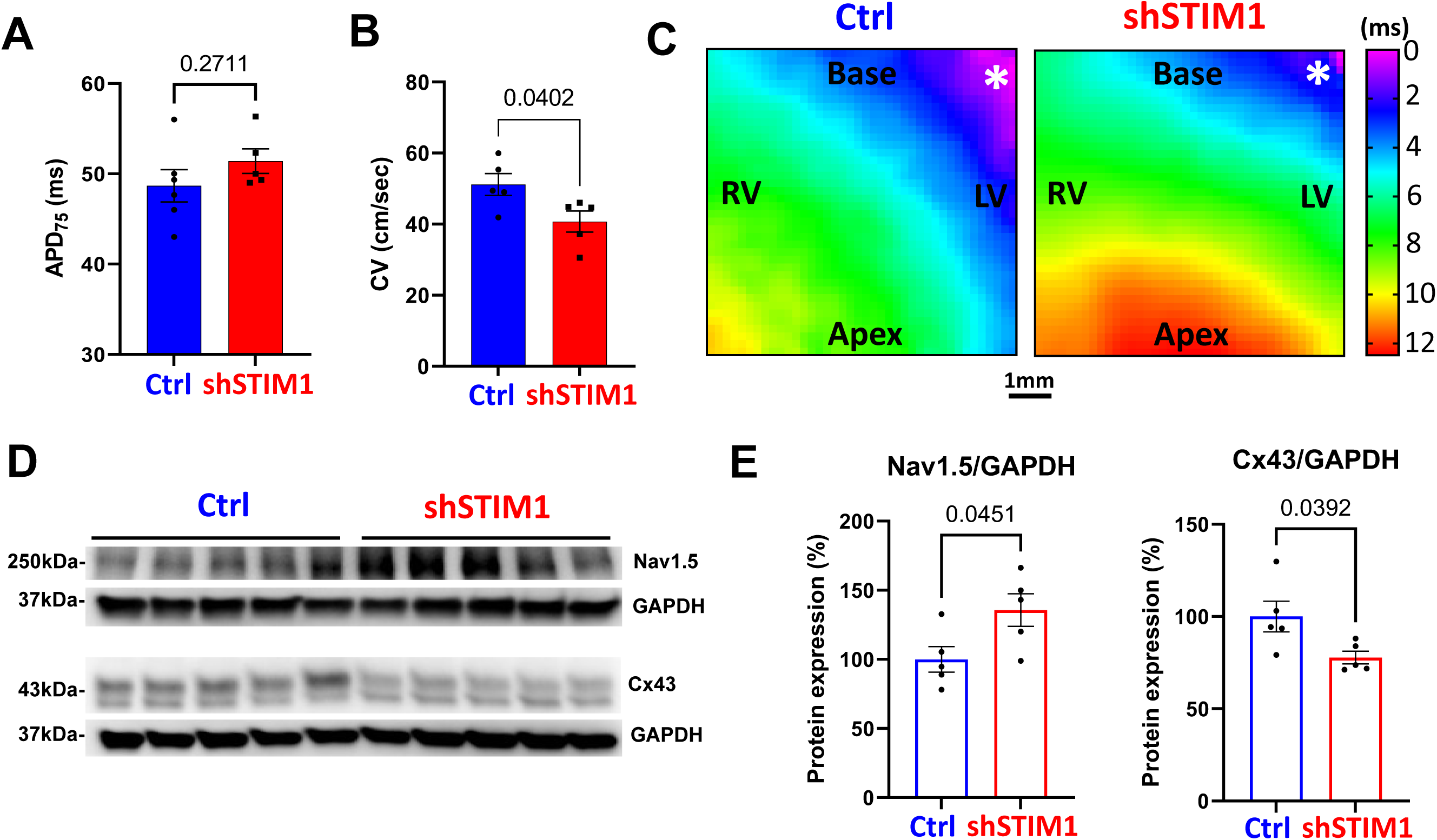
Electrophysiological effect of AAV9-mediated STIM1 silencing. **A.** *Ex-vivo* optical action potential mapping measurements showed similar action potential duration (APD_75_) in Ctrl (48.7±1.8ms) and shSTIM1 (51.4±1.4ms) hearts. N=6 Ctrl and 5 shSTIM1 hearts. Statistics: t-test. **B-C**. CV quantification and representative isochrone maps showing CV slowing in shSTIM1 hearts (40.7 ±3cm/sec vs 51.15 ±3.07cm/s in Ctrl hearts, p=0.04, PCL 140ms). N=5 hearts per group. Statistics: t-test. **D-E**. Representative immunoblots and quantification showing a significant increase in Nav1.5 and decrease in Cx43 protein expression in shSTIM1 hearts 4 weeks post gene transfer. N=5 hearts per group. Statistics: T-test.

We next challenged the hearts with rapid pacing. This protocol, which reliably induces Ca²⁺ overload, was expected to differentially stress hearts with STIM1 depletion. Representative AP recordings obtained during rapid pacing at a pacing cycle length (PCL) of 70 ms are shown in **Figure 3A**. In control hearts, even and odd beats displayed minimal variation in morphology, with no evidence of alternans behavior. In contrast, shSTIM1 hearts exhibited pronounced beat-to-beat differences in AP morphology and duration, characterized by alternating long and short APs, hallmarks of steady-state repolarization alternans and a well-established indicator of heightened arrhythmic vulnerability. Spatially, alternans extended across much of the mapped ventricular surface in shSTIM1 hearts (**Figure 3B**). Not only was the magnitude of alternans significantly greater in shSTIM1 compared with control hearts (**Figure 3C-E**), but the onset of alternans also occurred at slower heart rates, reflecting a clear leftward shift of the alternans threshold toward more physiological pacing frequencies (**Figure 3F**) indicative of higher vulnerability. Together, these findings indicate that STIM1 downregulation was associated with enhanced susceptibility to rate-dependent repolarization instability, a key substrate for ventricular arrhythmogenesis. Indeed, as shown in **Figure 4A**, further acceleration of the pacing rate (down to a PCL of 50 ms) elicited pronounced pro-arrhythmic behavior in the vast majority of shSTIM1 hearts (9/10), but rarely in controls (2/10; *p* < 0.0055; **Figure 4B**). Specifically, shSTIM1 hearts displayed a heightened propensity for loss of 1:1 capture and/or the onset of sustained episodes of VT/VF (**Figure 4C**).

**FIGURE 3:**
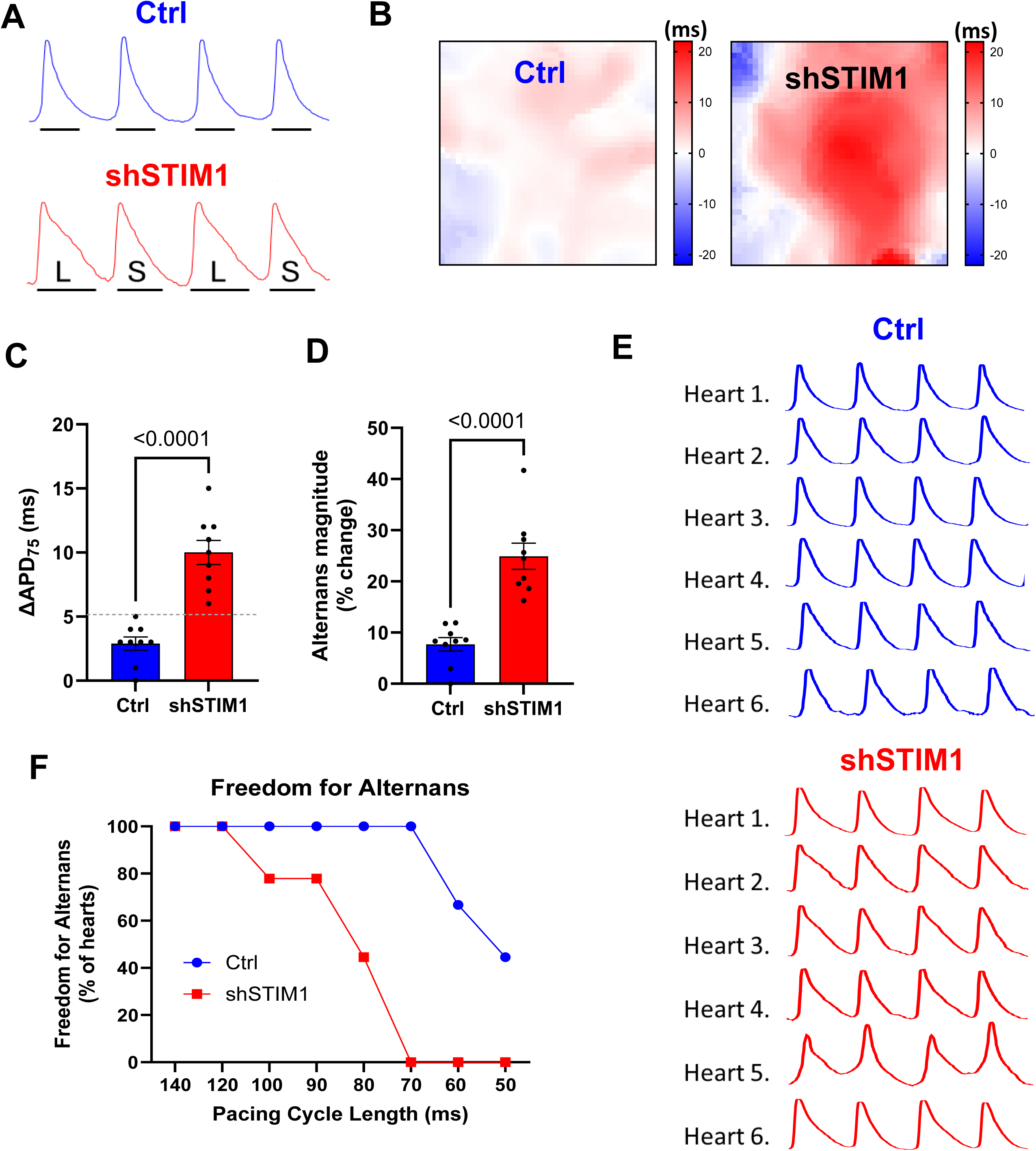
AAV9-mediated STIM1 silencing promotes pacing-induced alternans. **A**. Action potential (AP) traces recorded from representative Ctrl and shSTIM1 hearts during rapid pacing at PCL 70ms document presence of substantial beat to beat variability (alternans) in shSTIM1 but not in Ctrl hearts. **B**. Contour maps depicting the spatial distribution of ΔAPD_50_ in Ctrl and shSTIM1 hearts at pacing cycle length of 70ms. shSTIM1 but not Ctrl hearts exhibited AP alternans. **C**. Average ΔAPD_50_ measurement in Ctrl and shSTIM1 hearts measured at PCL70ms. N=9 hearts per group. Statistics: T-test. **D**. Percent change in magnitude of AP alternans measured as follows ((Long APD_50_-Short APD_50_)/Long APD_50_x100). N=9 hearts per group. Statistics: T-test. **E**. Representative action potential tracings from Ctrl and shSTIM1 hearts showing patterns of long-short AP alternans in shSTIM1 hearts. **F**. STIM1 silencing per se accelerates the development of AP alternans. The PCL at which AP alternans was first observed in shSTIM1 hearts was slower than that for Ctrl heart indicative of heightened propensity. Hearts were considered positive for presenting alternans when alternans magnitude was >or equal to 5ms between 2 consecutive action potentials. N=9 hearts per group.

**FIGURE 4:**
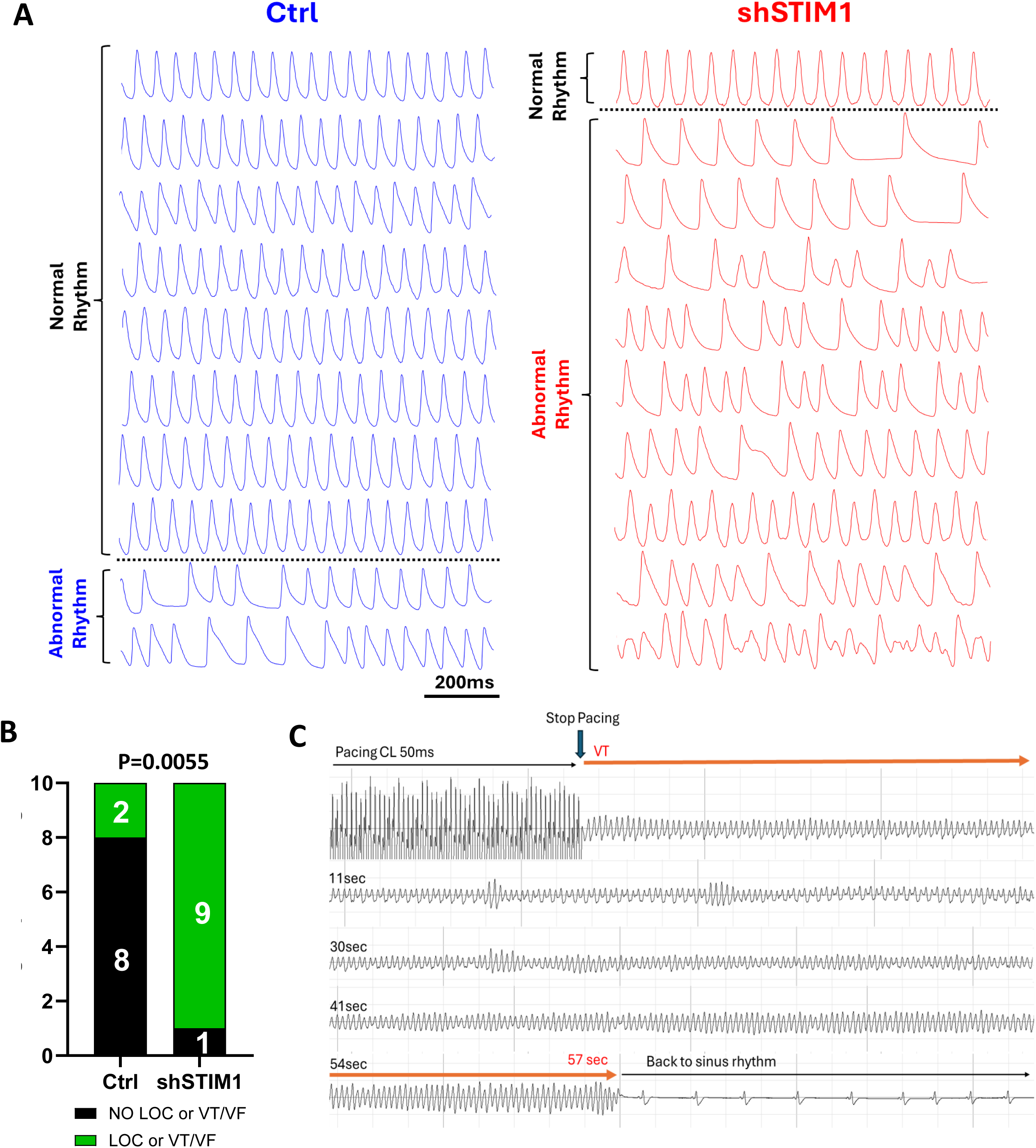
AAV9-mediated STIM1 silencing promotes pro-arrhythmic responses to heart rate elevation. **A**. Optical action potential traces of each of the 10 Ctrl and 10 shSTIM1 hearts showing arrhythmic activity marked by either loss of pacing capture or onset of VT in the vast majority of shSTIM1 but not Ctrl hearts in response to the pacing protocol which entailed pacing from PCL 100ms to 50ms in a Δ10ms decrements. **B**. Bar graph showing the number of hearts presenting with rapid pacing (up to 50ms PCL) induced arrhythmic behavior indexed by either loss of capture (<LOC) or onset of sustained VT/VF in all Ctrl and shSTIM1 hearts. Statistics: Fisher’s exact test. **C**. Volume-conducted ECG recording from an *ex-vivo* perfused shSTIM1 heart during showing onset of sustained VT/VF.

Next, we performed electron microscopy on Ctrl and shSTIM1 hearts. These studies revealed profound disruption of mitochondrial architecture in shSTIM1 hearts. Specifically, the mitochondrial network in shSTIM1 hearts exhibited abundance of rounded, fragmented, and disjointed organelles (**Figure 5A**). Quantitative analysis showed an approximately 20% reduction in mitochondrial perimeter and a 30% decrease in mitochondrial area relative to controls (**Figure 5B**). These ultrastructural abnormalities were accompanied by remodeling of mitochondrial dynamics proteins. Specifically, shSTIM1 hearts exhibited a 55% increase in phosphorylated DRP1 at Ser616 (pS616-DRP1/total DRP1 ratio: 1.55 ± 0.11 vs. 1.00 ± 0.09 in Ctrl, *p* = 0.0057), along with a more modest 21% increase in pS637-DRP1 (1.21 ± 0.04 vs. 1.00 ± 0.03 in Ctrl) (**Figure 5C-D**). Expression of the mitochondrial fusion protein OPA1 was downregulated by 15.4% (84.6 ± 2.4% vs. 100 ± 5.8% in Ctrl, *p* < 0.05), with no significant changes in MFN1 or MFN2 levels (**Figure 5E-F**).

**FIGURE 5:**
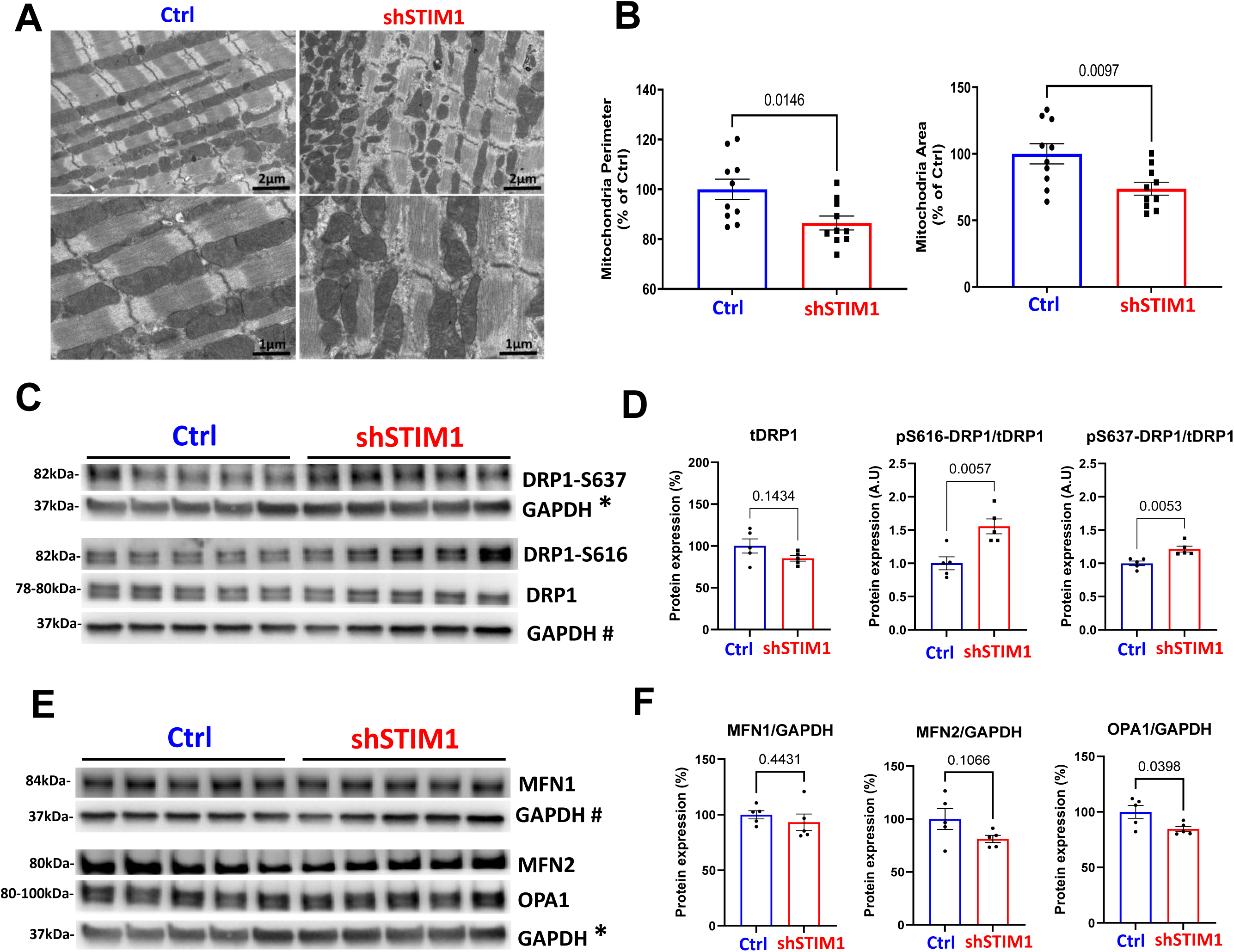
Cardiomyocyte STIM1 knockdown alters mitochondrial network ultrastructure and triggers mitochondrial fission. **A**. Transmission electron microscopy images from the LV of Ctrl and shSTIM1 hearts 4 weeks post gene transfer showing differences in mitochondrial network ultrastructure and organization. **B**. Mitochondrial dimensions (perimeters and areas) were quantified. **C-D**. Representative immunoblots and densitometric quantifications of total and phosphorylated DRP1 at Ser 637 and Ser 616 in tissue lysates from Ctrl and shSTIM1 hearts, 4 weeks post gene transfer. DRP1 protein levels were normalized to GAPDH and pDRP1 proteins were normalized to total DRP1 levels. shSTIM1 hearts showed a 55% and 21% increase in pS616/tDRP1 ratio (1.55±0.11 vs 1±0.09 in Ctrl) and p637/tDRP1 ratio (1.21±0.04 vs 1±0.03 in Ctrl), respectively versus Ctrl hearts. N=5 hearts per group was used. **E-F**. Representative immunoblots and densitometric quantifications of the major mitochondrial fission/fusion proteins, Mitofusin-1 (MFN1), Mitofusin-2 (MFN2) and OPA1 normalized to GAPDH in lysates from Ctrl and shSTIM1 hearts 4 weeks post gene transfer. While MFN1 and MFN2 protein levels were unchanged, OPA1 was decreased by 15.4% (84.6±2.4% vs 100±5.8% in Ctrl hearts). N=5 hearts per group. Statistics: t-test. **GAPDH\*** and **GAPDH#** mean that the same GAPDH blot was used to normalize the expression of multiple proteins (as indicated in Panels C and E) after stripping the membrane and blotting for those proteins.

Because both mitochondrial dynamics and STIM1 function are regulated by AMPK-dependent signaling,^26^ we next examined whether alterations in this pathway accompanied the observed mitochondrial remodeling. Despite comparable levels of total AMPK levels between groups (**Figure 6A&B**), shSTIM1 hearts exhibited marked suppression of AMPK downstream signaling. Specifically, phospho-to-total ACC ratio was reduced by 30% (0.69 ± 0.05 vs. 1.00 ± 0.04, p = 0.0014; **Figure 6A&C**), consistent with elevated malonyl-CoA synthesis and reduced mitochondrial fatty acid uptake for oxidation. Similarly, phospho-to-total Raptor levels were decreased by 33% (0.67 ± 0.06 vs. 1.00 ± 0.04, p = 0.002; **Figure 6A&D**), further indicating impaired AMPK signaling by STIM1 knockdown.

**FIGURE 6:**
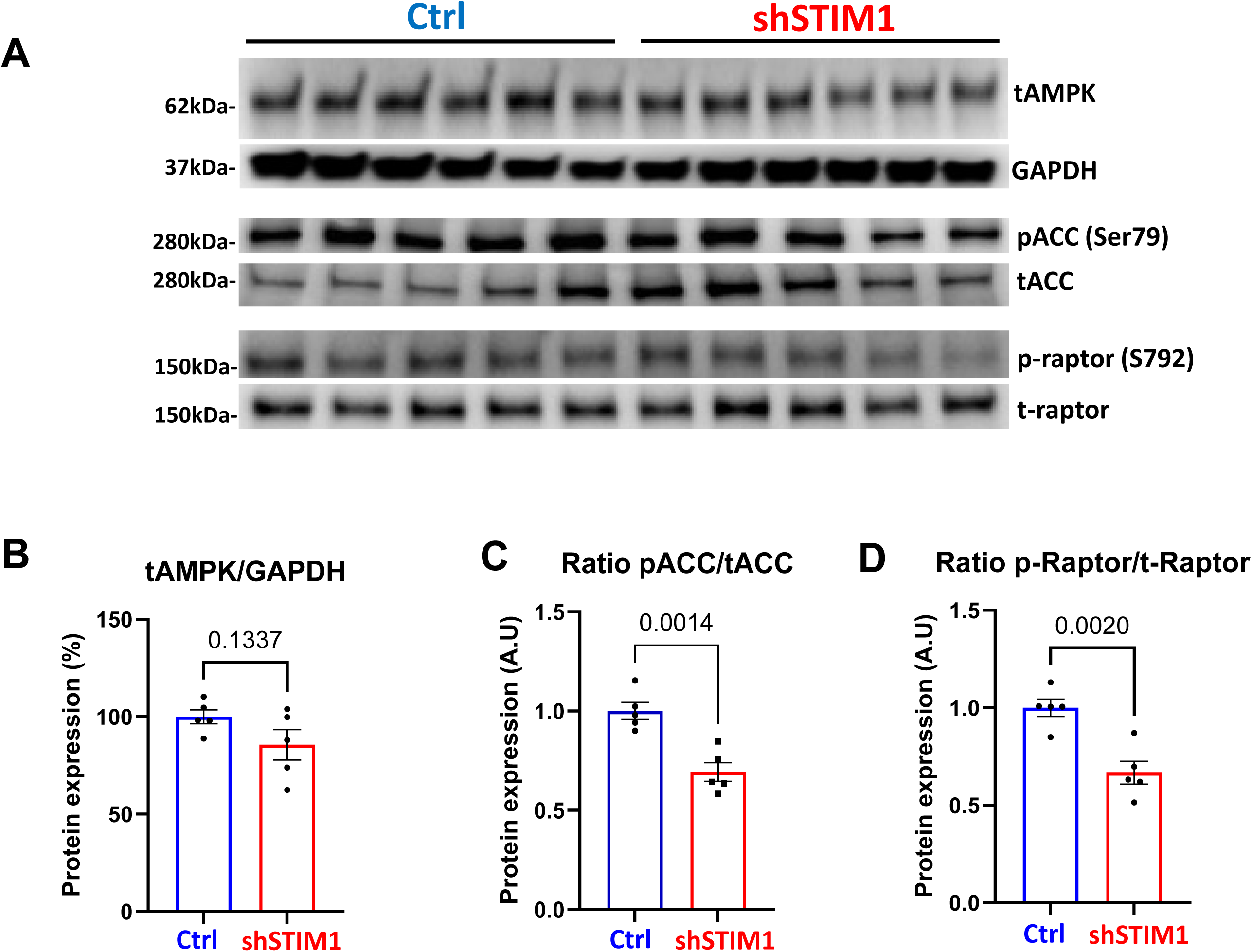
Cardiomyocyte STIM1 downregulation impairs AMPK downstream signaling. **A.** Representative immunoblots of AMPK, ACC, phospho-ACC-Ser79, raptor and phospho-raptor-Ser792 from cardiac lysates of Ctrl and shSTIM1 hearts 4 weeks post gene transfer. **B-D**. Densitometric quantifications of total AMPK and phosphorylated-to-total ratios for ACC and Raptor. N=5 hearts per group. Statistics: T-test.

Because STIM1 downregulation caused mitochondrial network remodeling and altered metabolic signaling, most notably reflected by reduced p-ACC levels, we hypothesized that these animals would be particularly susceptible to ischemia/reperfusion (I/R) injury. To test this, another cohort of mice was systemically administered either AAV9-shSTIM1 or a control vector. Four weeks later, animals underwent *in vivo* left coronary artery (LCA) occlusion for 30 minutes followed by reperfusion to induce MI. One week later, cardiac function was assessed by echocardiography, and hearts were subsequently harvested for high-resolution optical mapping of action potentials (**Figure 7A-B**). As shown in **Figure 7C**, STIM1 knockdown markedly exacerbated the response to MI as reflected by a much more pronounced decline in fractional shortening, significant thinning of the ventricular walls (indexed by IVSd and LVPWd), and substantial LV cavity dilation (LVIDd). On average, control hearts exhibited ∼15% reduction in fractional shortening one week after MI, whereas shSTIM1 hearts showed a decline exceeding 40% (**Figure 7D**, p < 0.0001). Consistent with these findings, MI-induced increases in LVIDd and decreases in LVPWd were also more pronounced in shSTIM1 hearts relative to controls, reflecting accelerated progression toward end-stage heart failure (**Figure 7D**).

**FIGURE 7:**
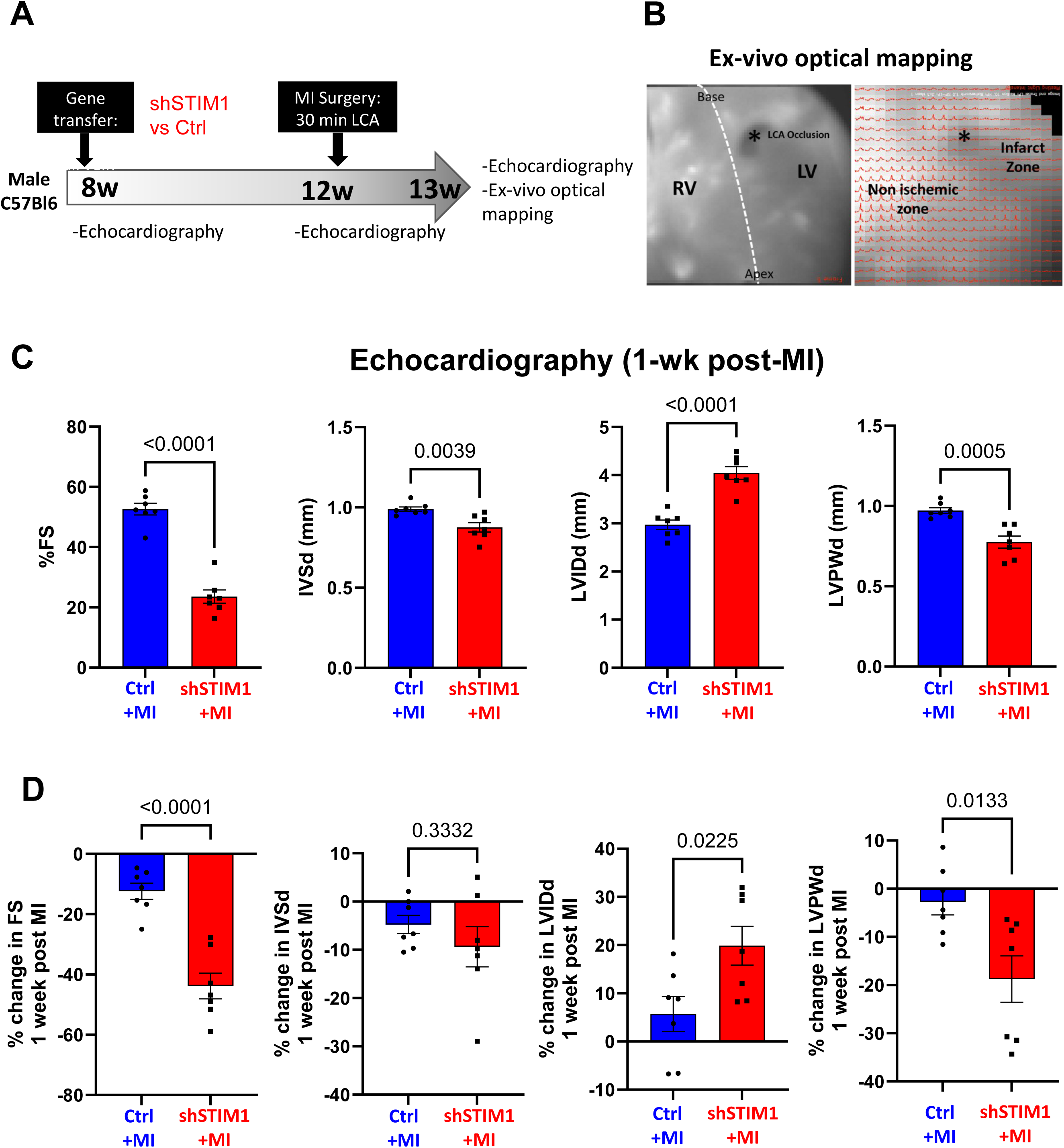
Post-MI mechanical and structural remodeling are exacerbated by AAV9-mediated STIM1 silencing. **A.** Schematic illustrating the experimental timeline to assess the impact of cardiomyocyte STIM1 downregulation on post-MI cardiac function. After baseline echocardiographic measurements, 8 weeks old C57Bl6/J male mice underwent AAV9-mediated shSTIM1 gene transfer via a tail vein injection with 1×10^11^ vg. Control (Ctrl) mice received either an AAV9 carrying a short hairpin against luciferase (shLuci) or PBS injection. Four weeks post gene transfer (at 12 weeks of age), mice underwent echocardiographic measurements. They were then randomized for cardiac MI operation by surgical LCA occlusion for 30min followed by reperfusion. One-week post-MI surgery (at 13 weeks of age), mice underwent a third round of echocardiographic measurements and were then sacrificed for ex vivo EP measurements. **B**. Ex-vivo optical mapping of an infarcted heart. Anatomical image of the mapped epicardial surface (left) and corresponding optical action potentials (AP) in a representative post-MI heart. **C**. Echocardiographic assessment of LV remodeling induced by AAV9-mediated STIM1 silencing 1-week post-MI (13-weeks old) in Ctrl and shSTIM1 mice. Parameters measured include fractional shortening (FS), interventricular septum thickness (IVS), LV internal diameter (LVID) and LV posterior wall thickness (LVPW) during diastole (d). **D**. Percent change in FS, IVSd, LVIDd and LVPWd at 1w post MI (13w old) compared to prior-MI (12w old). N=7 AAV9.shSTIM1 mice and N=7 control mice were used for this analysis. Statistics: T-test.

To assess the EP substrate underlying these changes, high-resolution optical AP mapping was performed in post-MI hearts (**Figure 7B**). This analysis revealed reduction in CV in post-MI shSTIM1 hearts compared with post-MI controls across a wide range of PCLs (**Figure 8A-B**). Moreover, post-MI shSTIM1 hearts exhibited a trend toward shorter APD₇₅ (by −31.5%), driven by localized remodeling within the peri-infarct zone (**Figure 8C-E**). To distinguish between EP changes driven by STIM1 per se from those requiring the combined effects of STIM1 loss and MI, we compared the impact of AAV9-mediated STIM1 knockdown under both non-MI and MI conditions. As shown in **Figure 8F-G**, STIM1 downregulation produced a comparable ∼20% reduction in CV in both settings relative to their respective controls. In contrast, while STIM1 knockdown had minimal influence on APD under non-MI conditions (a non-significant 5.6% prolongation vs. Ctrl), it caused a marked 26.5% APD shortening following MI, indicating that the effects of STIM1 loss on repolarization are strongly dependent on disease state.

**FIGURE 8:**
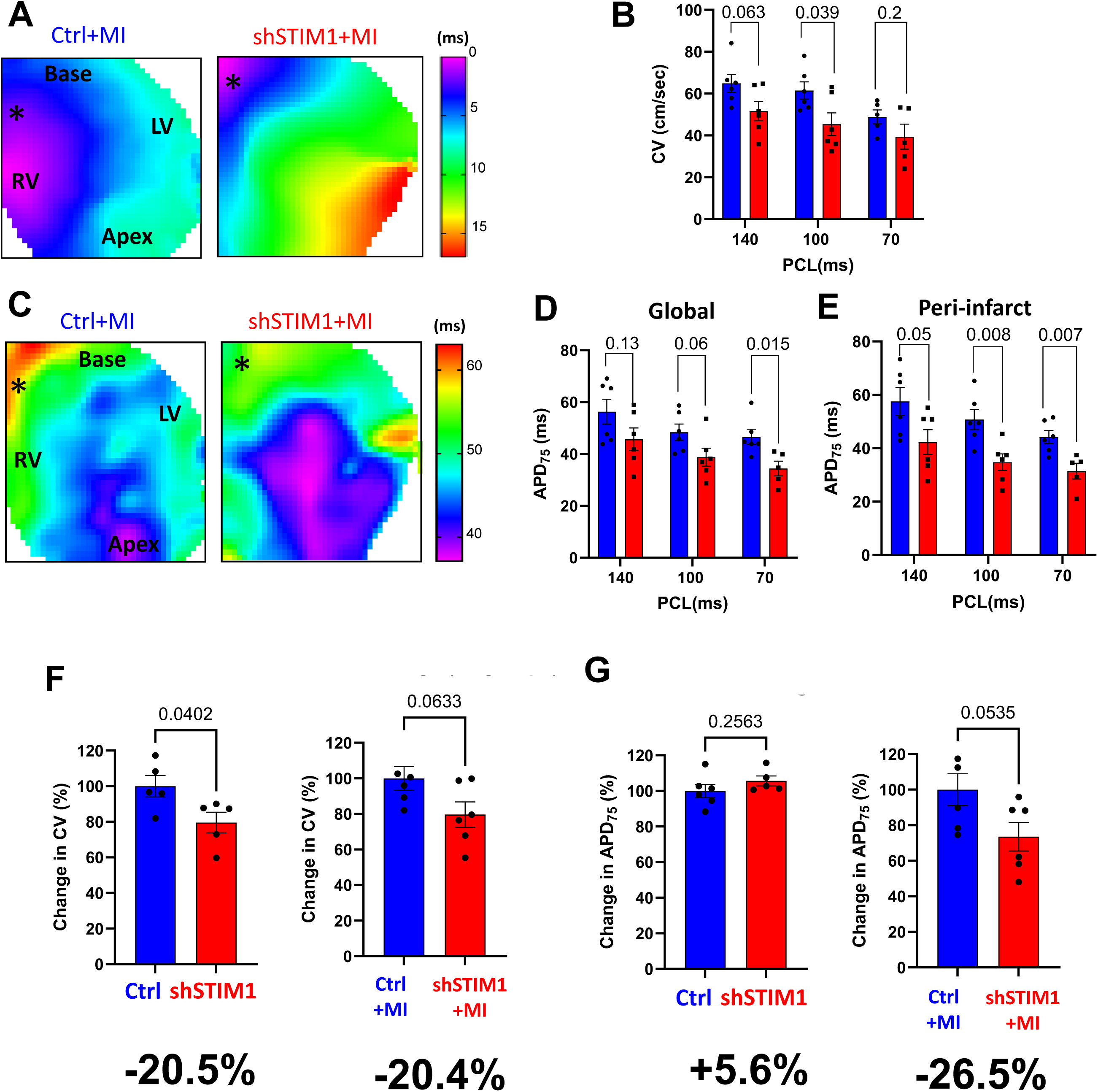
Cardiomyocyte STIM1 depletion exacerbates post-MI electrical remodeling. **A.** Representative depolarization maps depicting the spread of activation across the ventricular epicardium of post-MI Ctrl and post-MI shSTIM1 hearts. **B.** Average and distribution of CV of post-MI Ctrl (Ctrl+MI) and shSTIM1 (shSTIM1+MI) hearts measured at PCL 140, 100 and 70ms. N=6 Ctrl+MI and N=6 shSTIM1+MI hearts. Statistics: T-test between both groups at each PCL. **C.** Representative APD contour maps measured from a post-MI control (Ctrl+MI) and shSTIM1 (shSTIM1+MI) heart. **D-E**. Quantification of APD_75_ on the entire epicardial surface (global) and within the peri-infarct region at PCL of 140, 100 and 70ms. N=6 Ctrl+MI and N=6 shSTIM1+MI hearts. Statistics: T-Test between both groups at each PCL. **F**. Percent change in CV in the shSTIM1 hearts compared to Ctrl hearts prior to MI (Left graph) and 1-week post-MI (Right graph) showing a similar CV slowing of −20.5% and −20.4% respectively. Statistics: T-test. **G**. Percent change in APD_75_ for in shSTIM1 hearts compared to Ctrl hearts prior to MI (Left graph) and 1-week post-MI (Right graph) showing the appearance of APD shortening in shSTIM1 hearts by - 26.5% post MI. Statistics: T-test.

Finally, we assessed the response of all groups to rapid pacing (**Figure 9A**). Spatially discordant AP alternans emerged in 67% post-MI hearts with STIM1 downregulation but were rarely seen in hearts with either condition alone (**Figure 9B-C**). This indicates that both LV dysfunction and STIM1 loss are jointly required to trigger this complex, highly arrhythmogenic behavior. Overall, these findings reveal a synergistic interaction between MI and STIM1 downregulation in promoting the emergence of spatially discordant alternans.

**FIGURE 9:**
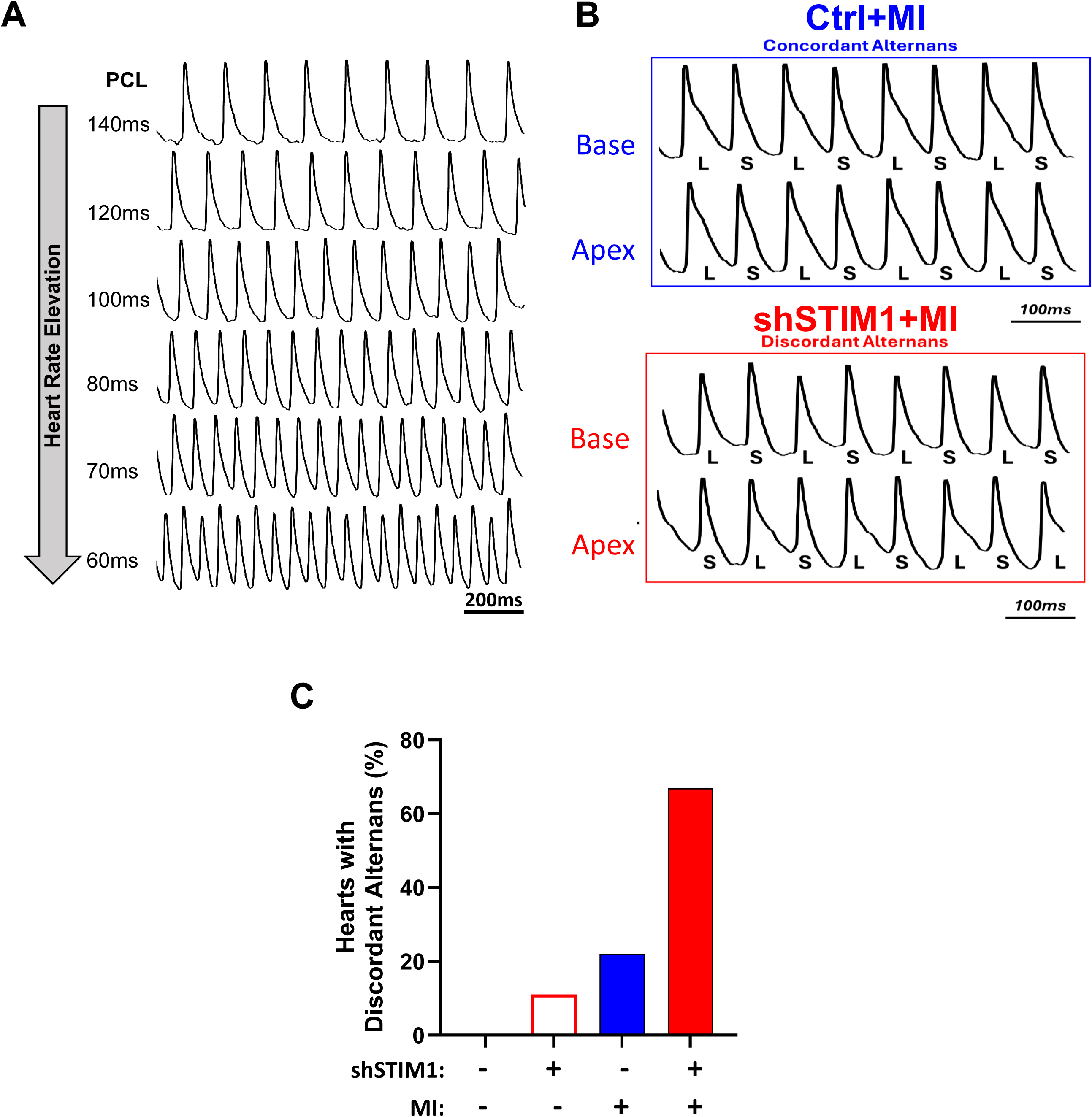
The combination of AAV9-mediated STIM1 silencing and MI induced LV failure triggers discordant alternans behavior. **A**. Representative AP traces recorded from a post-MI shSTIM1 hearts (shSTIM1+MI) showing the emergence of beat-to-beat AP alternans at elevated pacing rates (PCL shortening). **B**. Action potential traces recorded at PCL of 70ms. Post-MI shSTIM1 hearts exhibited more severe alternans patterns that were spatially discordant compared to their post-MI Ctrl counterparts. **C.** Percentage of hearts exhibiting spatially discordant AP alternans in the 4 groups: control, shSTIM1 only, MI only, and combine MI with shSTIM1.

## DISCUSSION

The primary objective of this study was to determine how myocardial STIM1 downregulation alters electromechanical function and modulates the heart’s response to myocardial infarction. To achieve this, we employed an AAV9-mediated gene transfer strategy designed to overcome key limitations of the tamoxifen-inducible MerCreMer model, in which abrupt STIM1 depletion results in rapid mortality (>50% within eight days).^13^ This high lethality confounds interpretation by making it difficult to distinguish electrophysiological changes that arise directly from STIM1 loss versus those secondary to severe hemodynamic instability. In contrast, AAV9-mediated knockdown enabled a gradual and sustained reduction of STIM1 expression in the adult heart without precipitating acute heart failure. This approach thus circumvented both the acute lethality associated with inducible deletion and the developmental reprogramming inherent to constitutive knockout models. Using this platform, we were able to directly assess the electrophysiological consequences of STIM1 downregulation, examine associated alterations in mitochondrial structure and metabolic signaling, and determine how these baseline perturbations influence post-MI remodeling, particularly in light of prior findings that smooth muscle STIM1 depletion confers protection against ischemia/reperfusion injury.^22^

### Myocardial STIM1 downregulation alters electro-mechanical function and promotes EP instability

Cardiac function was assessed at baseline (time of gene delivery) and again four weeks later to evaluate the direct impact of reduced STIM1 expression on myocardial performance. Hearts receiving AAV9-shSTIM1 exhibited a modest but consistent decline in left ventricular contractile function accompanied by chamber remodeling. These findings reinforce and extend prior observations by demonstrating that diminished STIM1 expression alone is sufficient to impair cardiac performance, even in the absence of external stressors such as pressure overload.

Notably, AAV9-mediated STIM1 knockdown also produced marked conduction slowing associated with reduced Cx43 expression, unveiling a previously unrecognized functional link between STIM1, an ER/SR calcium sensor and Cx43, the principal gap junctional protein governing electrical coupling and impulse propagation. This novel association highlights a potential cross-talk between calcium-sensing pathways and intercellular communication in the adult heart. Whether this relationship arises from a direct molecular interaction or reflects indirect metabolic perturbations that influence Cx43 synthesis, trafficking, or turnover remains an important question for future investigation.

Furthermore, STIM1 downregulation alone increased susceptibility to proarrhythmic behavior, evidenced by the early onset of action potential alternans. These findings parallel our previous results in tamoxifen-inducible, cardiac-specific STIM1 knockdown mice^13^ and reinforce the concept that loss of myocardial STIM1 intrinsically disrupts baseline electrophysiological homeostasis. Even in the absence of significant LV dysfunction or heart failure, diminished STIM1 expression creates a vulnerable substrate.

### Myocardial STIM1 downregulation causes remodeling of mitochondrial ultrastructure and metabolic signaling

To elucidate the mechanisms by which STIM1 loss disrupts cardiac EP function, we next focused on its impact on mitochondrial dynamics and metabolic signaling. Indeed, we found that mechanical dysfunction following AAV9-mediated myocardial STIM1 downregulation coincided with pronounced remodeling of the mitochondrial network, underscoring the tight coupling between STIM1-mediated Ca²⁺ sensing and mitochondrial integrity. Mitochondria displayed smaller, rounded profiles with reduced area and perimeter, features consistent with enhanced mitochondrial fission. These morphological changes were accompanied by increased phosphorylation of DRP1 at Ser616 and reduced OPA1 expression, confirming activation of the fission machinery. Similar mitochondrial alterations were previously described by Collins et al.^15^, who reported an increased number of smaller mitochondria associated with heightened fission in STIM1-deficient hearts.

Mitochondrial network architecture reflects the dynamic interplay among biogenesis, fusion, fission, and mitophagy—processes tightly coordinated by the master metabolic regulator AMPK ^26^. Given this hierarchical control, we next assessed AMPK downstream signaling by immunoblotting and observed significant reductions in the phospho-to-total ratios of both ACC and Raptor, indicating suppressed AMPK activity. The role of STIM1 in regulating metabolic signaling was previously examined by Collins et al^14^ using cardiac-restricted (Cr) STIM1 knockout mice. By 20 weeks of age, these mice exhibited profound metabolic reprogramming, including reduced glucose utilization, decreased GLUT4 expression, and diminished AMPK phosphorylation. Although fatty acid oxidation (FAO) rates remained unchanged, lipid and triglyceride accumulation increased in parallel with upregulation of FAO-related proteins, suggesting an imbalance between lipid uptake and oxidation. Whether these alterations reflected a direct consequence of STIM1 loss or secondary effects of chronic remodeling or developmental compensation, however, remained uncertain.

In our present study, AAV9-shSTIM1–mediated suppression of ACC phosphorylation in the adult heart reflects loss of AMPK-dependent inhibition of ACC, a shift expected to promote malonyl-CoA accumulation and consequent inhibition of mitochondrial FAO. This metabolic inflexibility restricts cardiac substrate utilization, favoring mitochondrial dysfunction, particularly under ischemic conditions.

### AAV9-mediated STIM1 downregulation exacerbates post-MI remodeling

Given STIM1’s central role in regulating intracellular Ca²⁺ homeostasis^12, 27, 28^ and its interactions with mitochondrial^29^ and metabolic signaling networks,^16, 30, 31^ we hypothesized that STIM1 serves as a critical determinant of post–MI remodeling, a process characterized by Ca²⁺ overload and mitochondrial dysfunction. Supporting this premise, STIM1 upregulation following MI has been linked to increased apoptosis.^32^ *In vitro*, hypoxia/reoxygenation in H9C2 cells induces a fourfold increase in STIM1 expression accompanied by a parallel rise in apoptosis, whereas siRNA-mediated STIM1 silencing markedly reduces cell death.^32^ In contrast, elegant work by Mali et al. demonstrated that selective deletion of STIM1 in smooth muscle cells protects against I/R injury by reducing infarct size and improving cardiac function.^22^ These divergent observations prompted us to test whether myocardial reduction of STIM1 levels using AAV9-mediated gene silencing might confer protection against or exacerbate adverse post-MI remodeling. Our findings show that myocardial STIM1 downregulation markedly aggravates post-MI structural and functional deterioration culminating in end-stage heart failure. This outcome extends prior findings that STIM1 downregulation accelerates the transition from compensated hypertrophy to decompensated heart failure under pressure overload stress.^11^ Collectively, these data indicate that myocardial STIM1 is indispensable for preserving electromechanical stability during stress and that its loss undermines key adaptive mechanisms essential for recovery after ischemic injury.

The contrasting outcomes of STIM1 depletion in the myocardium versus selectively in smooth muscle cells^22^ underscore the need to define the distinct, cell type–specific roles of STIM1 within the heart. These divergent effects further caution against the use of nonspecific STIM1 inhibitors, given that STIM1 suppression may elicit opposing outcomes across different cardiac cell populations involved in I/R injury. Together, our findings and those of Mali et al.^22^ emphasize the importance of delineating discrete, cell-specific STIM1-dependent mechanisms to enable the rational design of targeted therapeutic strategies that preserve protective STIM1 signaling in cardiomyocytes while mitigating its maladaptive effects in nonmyocyte populations.

### Direct and indirect effects of STIM1 on EP properties and the pro-arrhythmic substrate

By examining the effects of STIM1 downregulation in both non-MI and MI settings, we were able to distinguish electrophysiological alterations that arise directly from STIM1 loss independent of disease state from those that manifest only in the remodeled myocardium. STIM1 downregulation induced a comparable degree of CV slowing in both non-MI and MI hearts, suggesting a primary role for STIM1 in regulating myocardial conduction, likely mediated at least in part by reduced Cx43 expression. In contrast, APD shortening was observed only in the MI setting, indicating that this effect may depend on the altered pro-oxidant and/or metabolic milieu of the remodeled heart.

Finally, a major finding of this study is the identification of conditions that unmask spatially discordant AP alternans, a proarrhythmic behavior previously linked by our group and many others to the initiation of ventricular fibrillation. In the current work, the incidence of discordant alternans increased synergistically in the combined MI + shSTIM1 group. Whereas control, shSTIM1-only, and MI-only hearts exhibited low event rates, the combined group showed a marked rise in vulnerability. This indicates that the coexistence of the two factors creates a uniquely vulnerable electrophysiological substrate. We previously demonstrated the mechanistic significance of discordant alternans in an inducible, cardiomyocyte-specific STIM1 knockout model, where animals developed severe LV dysfunction (fractional shortening ≈20%). A similar degree of LV impairment was observed in the post-MI AAV9-shSTIM1 group, reinforcing the concept that severe mechanical dysfunction is a required feature in the emergence of discordant alternans.

## ABBREVIATIONS

STIM1: Stromal Interaction Molecule 1
SOCE: Store-operated calcium entry
SR/ER: Sarco/endoplasmic reticulum
VT/VF: Ventricular tachycardia/ventricular fibrillation
AAV9: Adeno-associated virus serotype 9
Ctrl: Control
MI: Myocardial infarction
Ctrl+MI: Post-MI control heart
shSTIM1+MI: Post-MI heart that underwent AAV9-mediated STIM1 silencing
PCL: Pacing cycle length
APD: Action potential duration
CV: Conduction velocity
Cx43: Connexin-43
FS: Fractional shortening
LVIDd: Left ventricular internal diameter during diastole
IVSd: Intraventricular septum thickness during diastole
LVPWd: Left ventricular posterior wall thickness during diastole

