## Supplementary figures and images for "Myocardial STIM1 deficiency triggers mitochondrial fission to impair electrophysiological function and exacerbate post-MI remodeling"

### Data Supplement 1

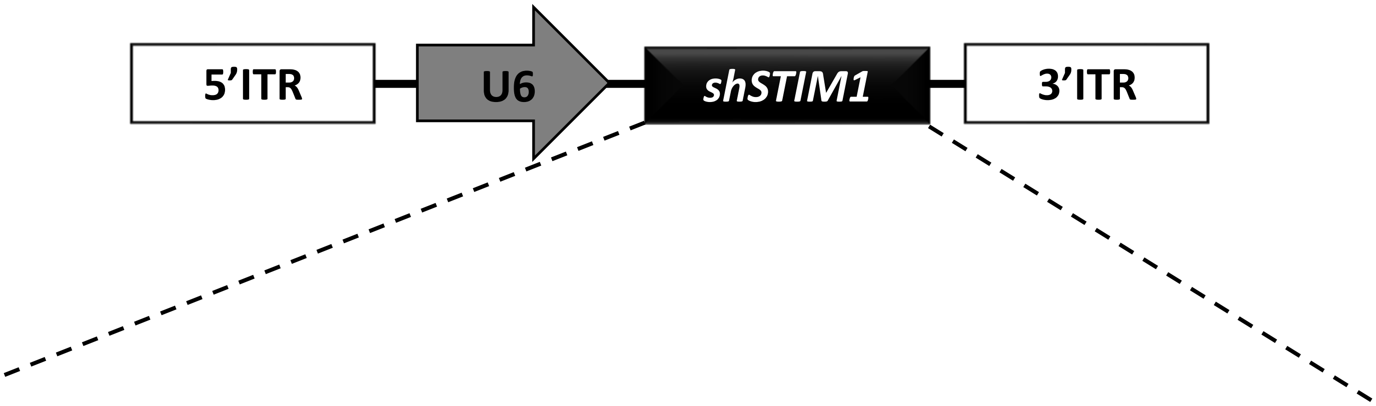


GGGAAGACCTCAATTACCATTCAAGAGATGGTAATTGAGGTCTTCCCTTTTTT
