## Supplementary material for "Myocardial STIM1 deficiency triggers mitochondrial fission to impair electrophysiological function and exacerbate post-MI remodeling": Data Supplement 2

### Slide 1
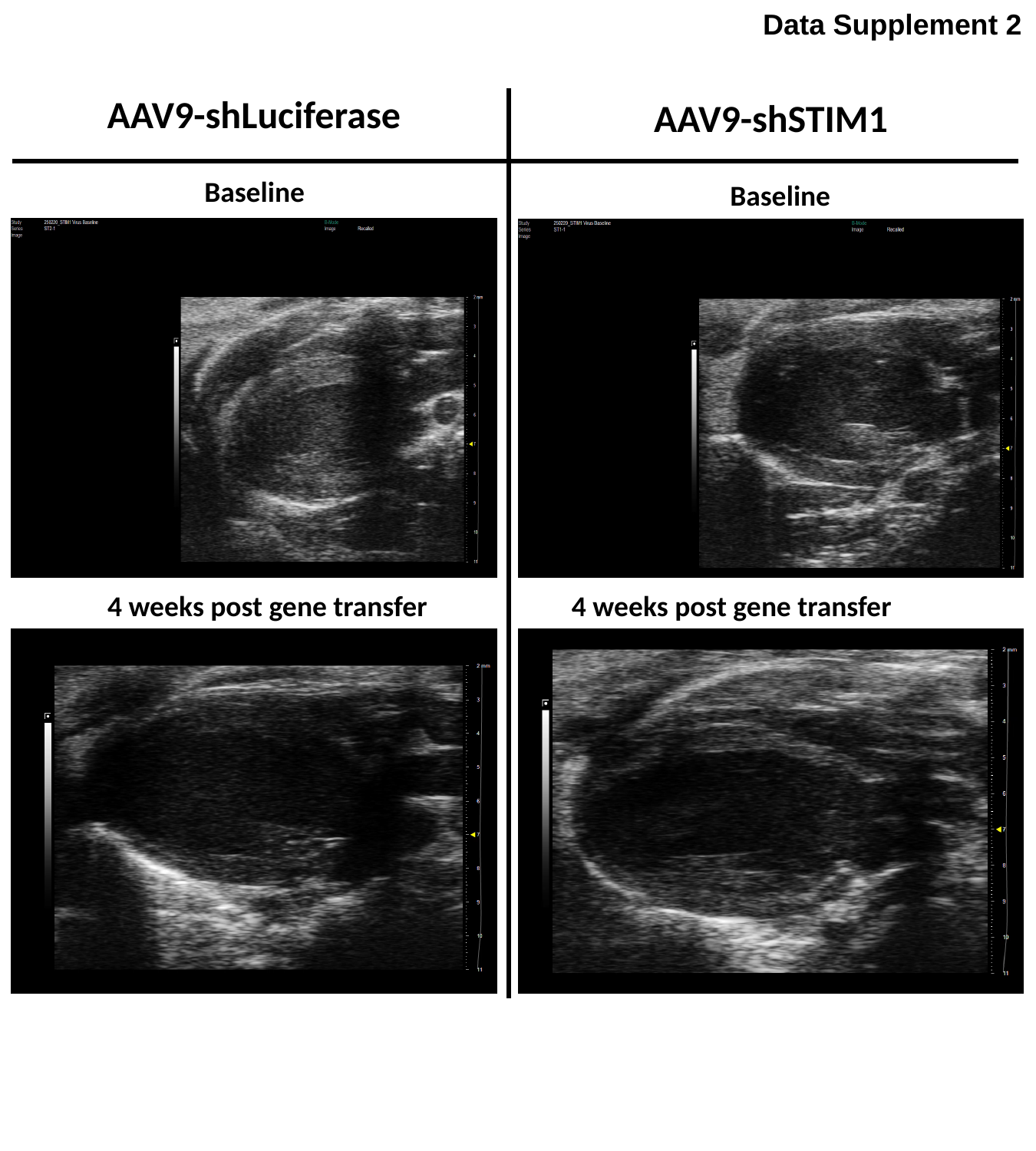

Data Supplement 2
AAV9-shLuciferase
AAV9-shSTIM1
Baseline
Baseline
4 weeks post gene transfer
4 weeks post gene transfer
